## Supplementary figures and images for "Coding of latent variables in sensory, parietal, and frontal cortices during closed-loop virtual navigation" (from another version of this preprint: version 2, 2022-05-13)

### Figure 1 Supplement 1

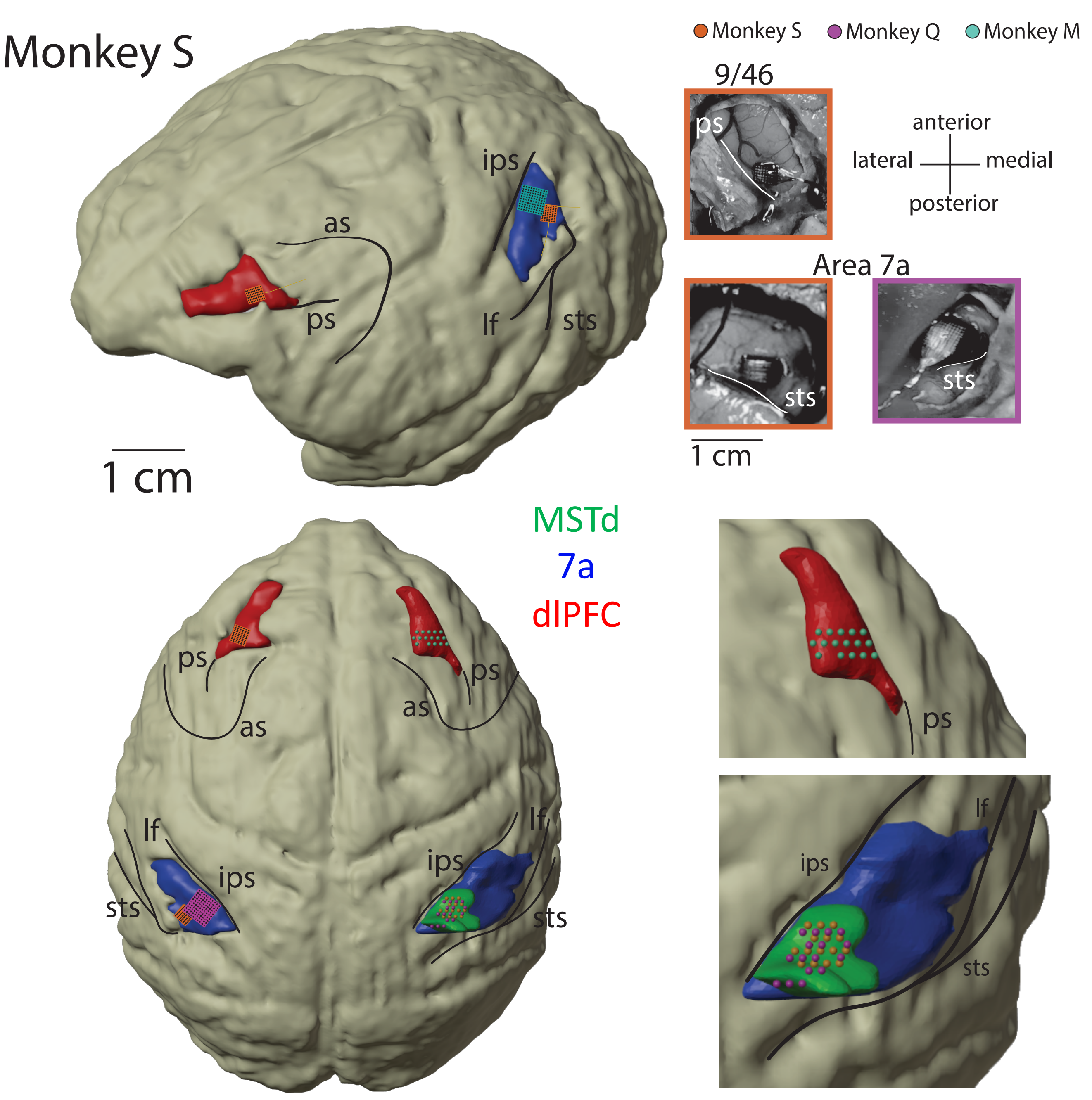

### Figure 2 Supplement 1

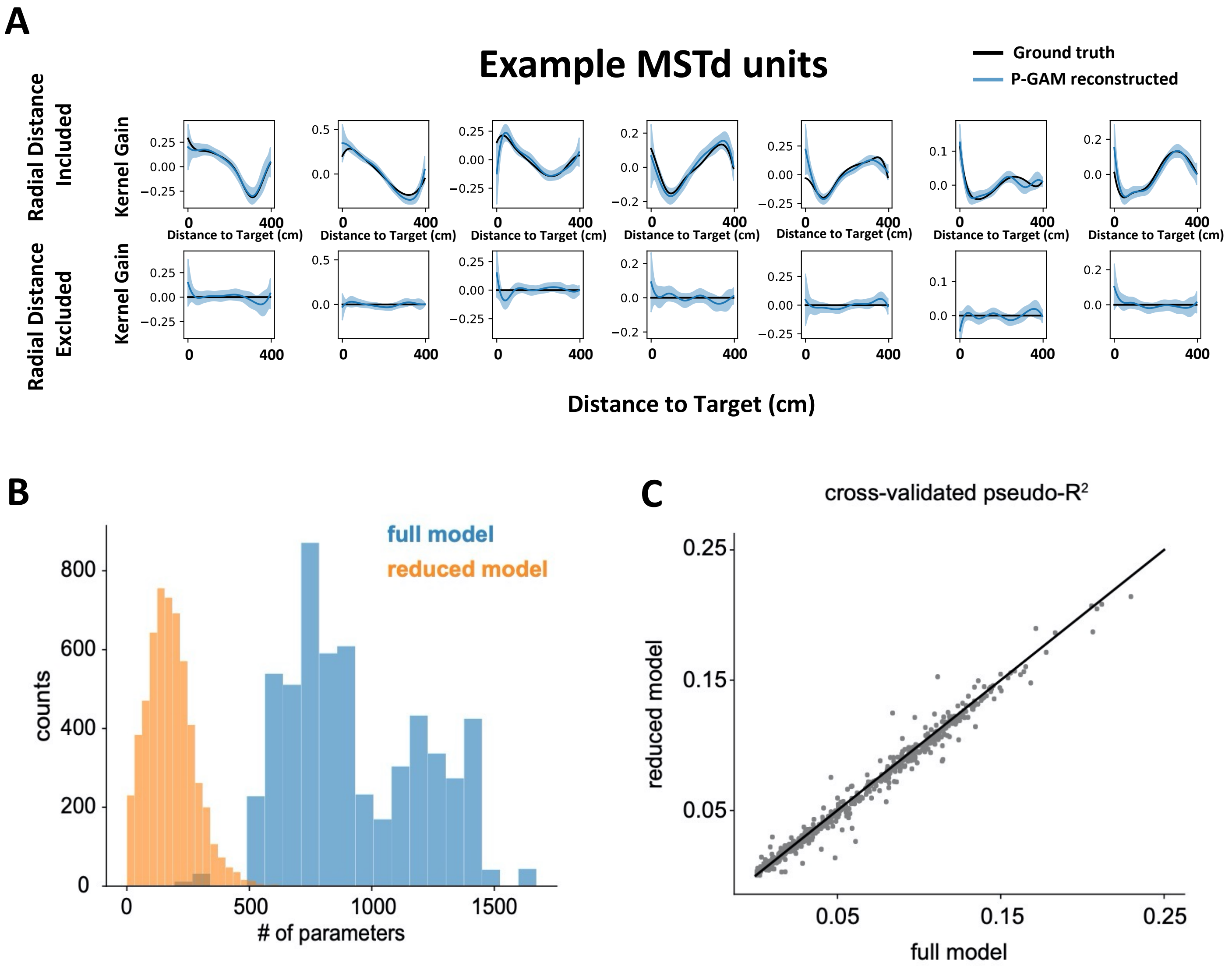

### Figure 2 Supplement 2

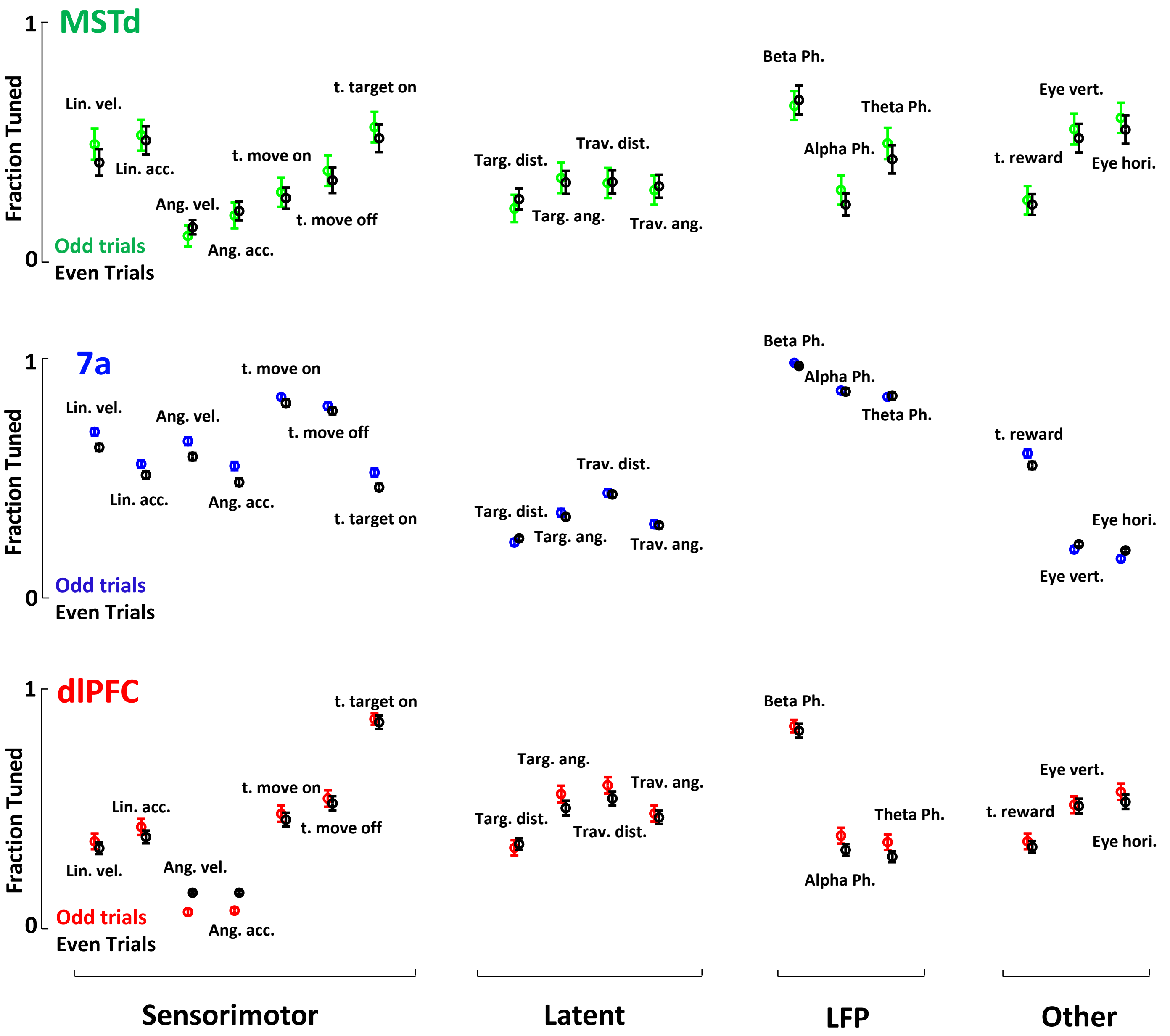

### Figure 3 Supplement 1

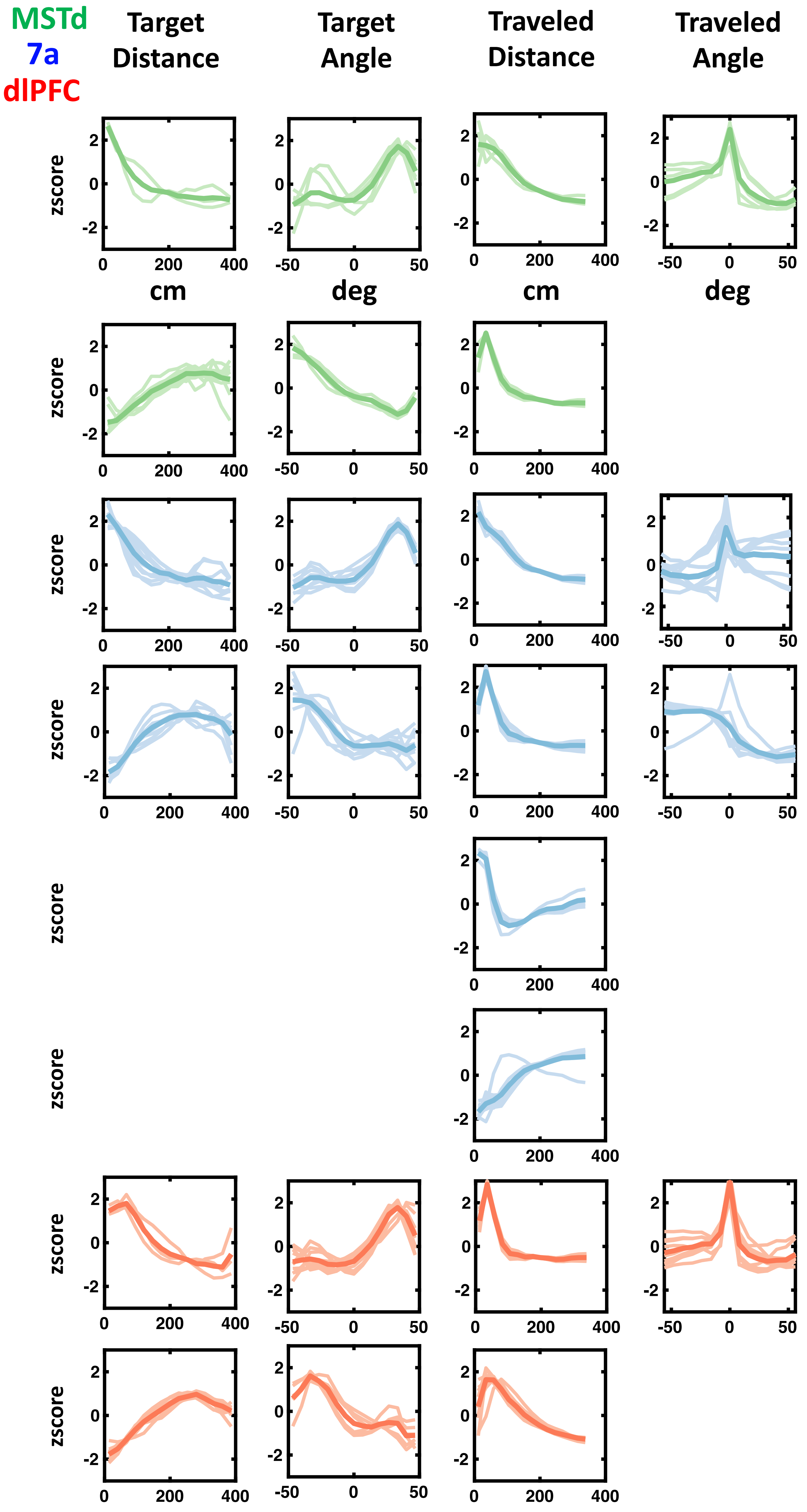

### Figure 3 Supplement 2

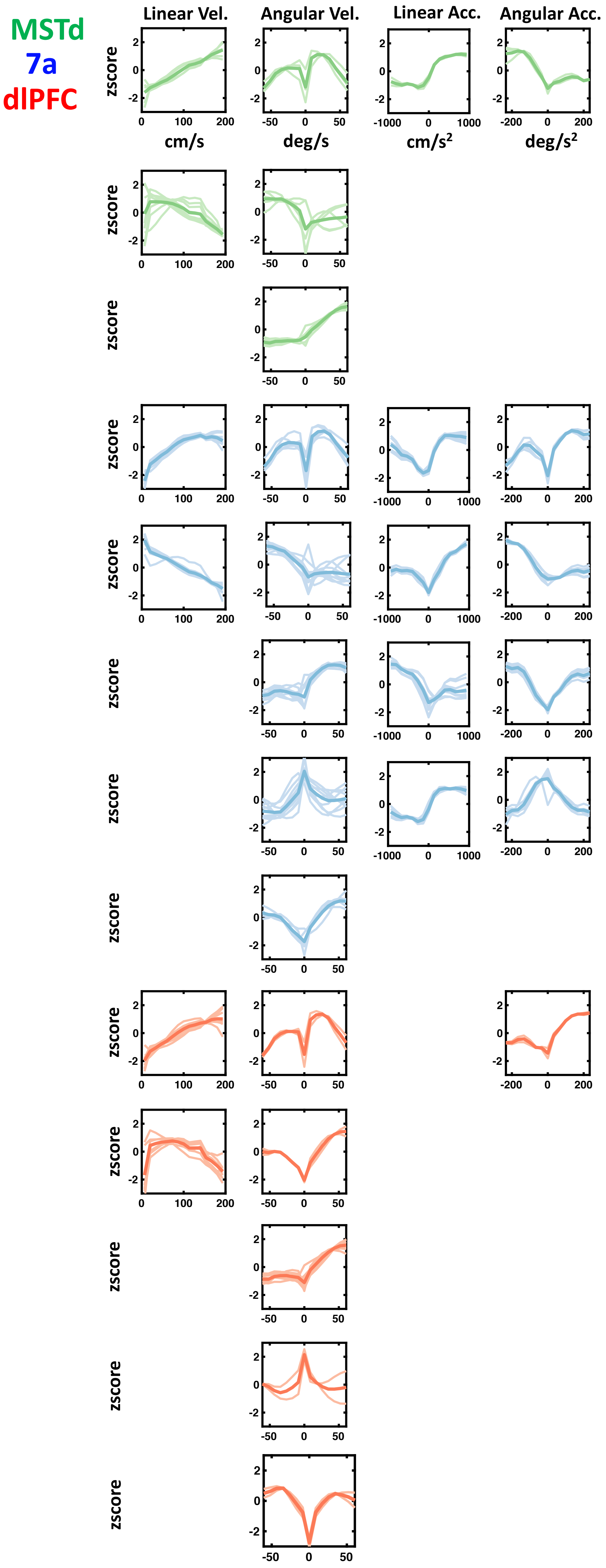

### Figure 3 Supplement 3

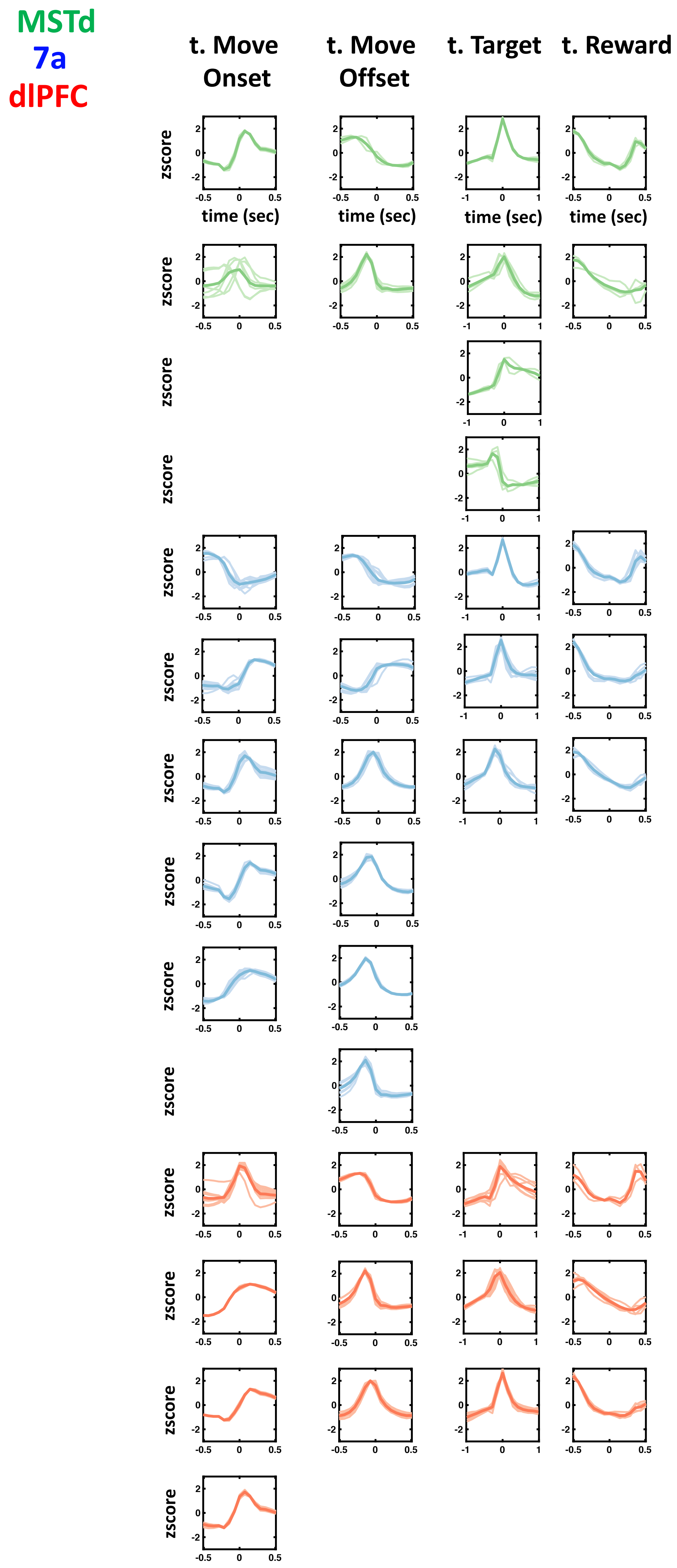

### Figure 3 Supplement 4

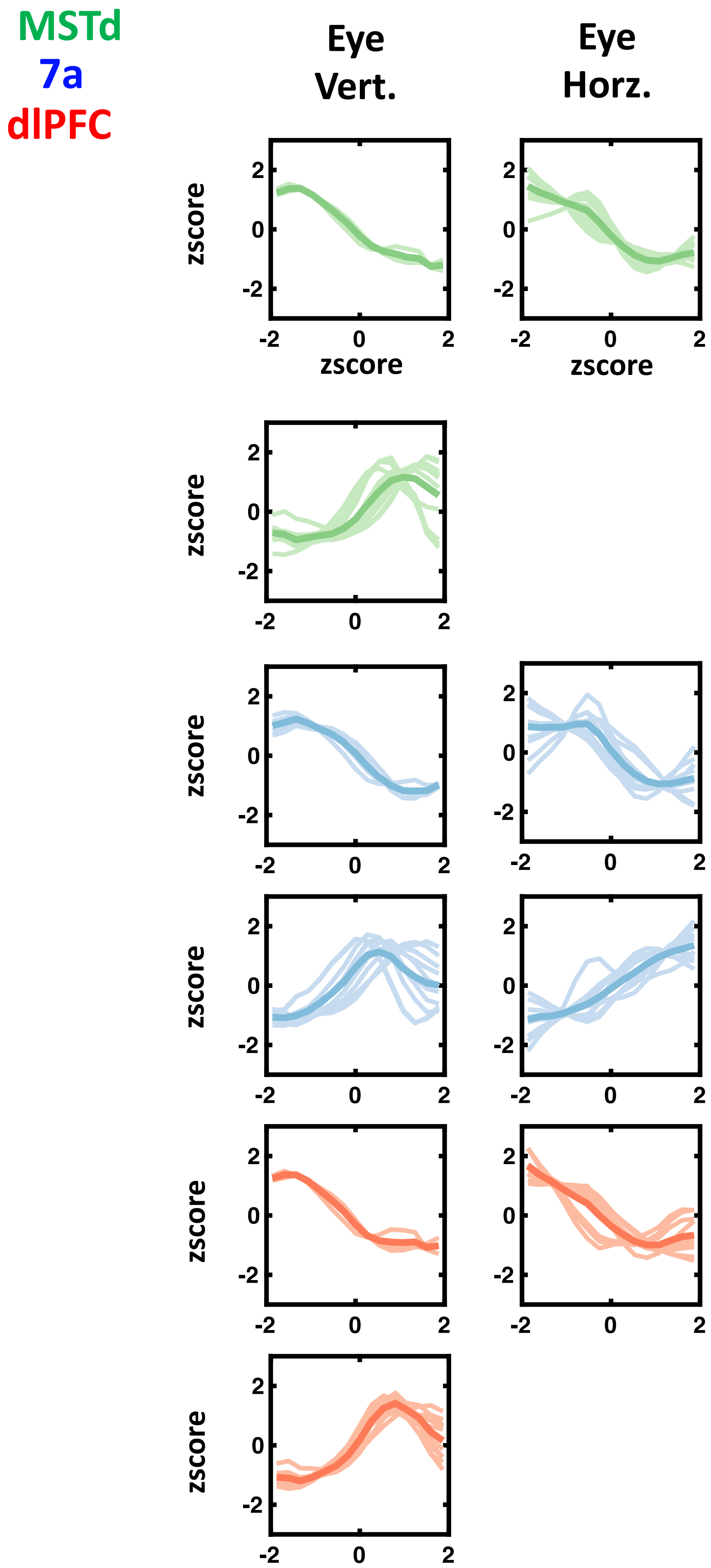

### Figure 4 Supplement 1

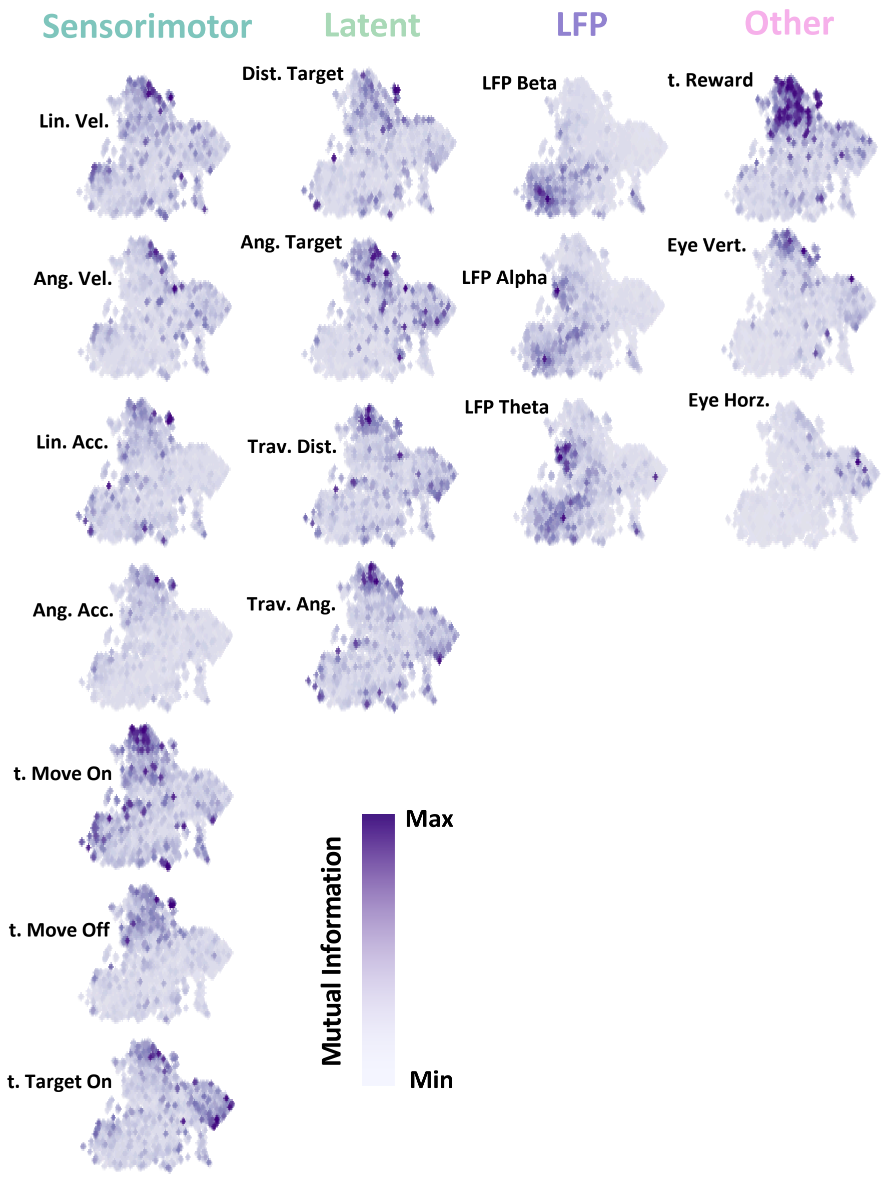

### Figure 4 Supplement 2

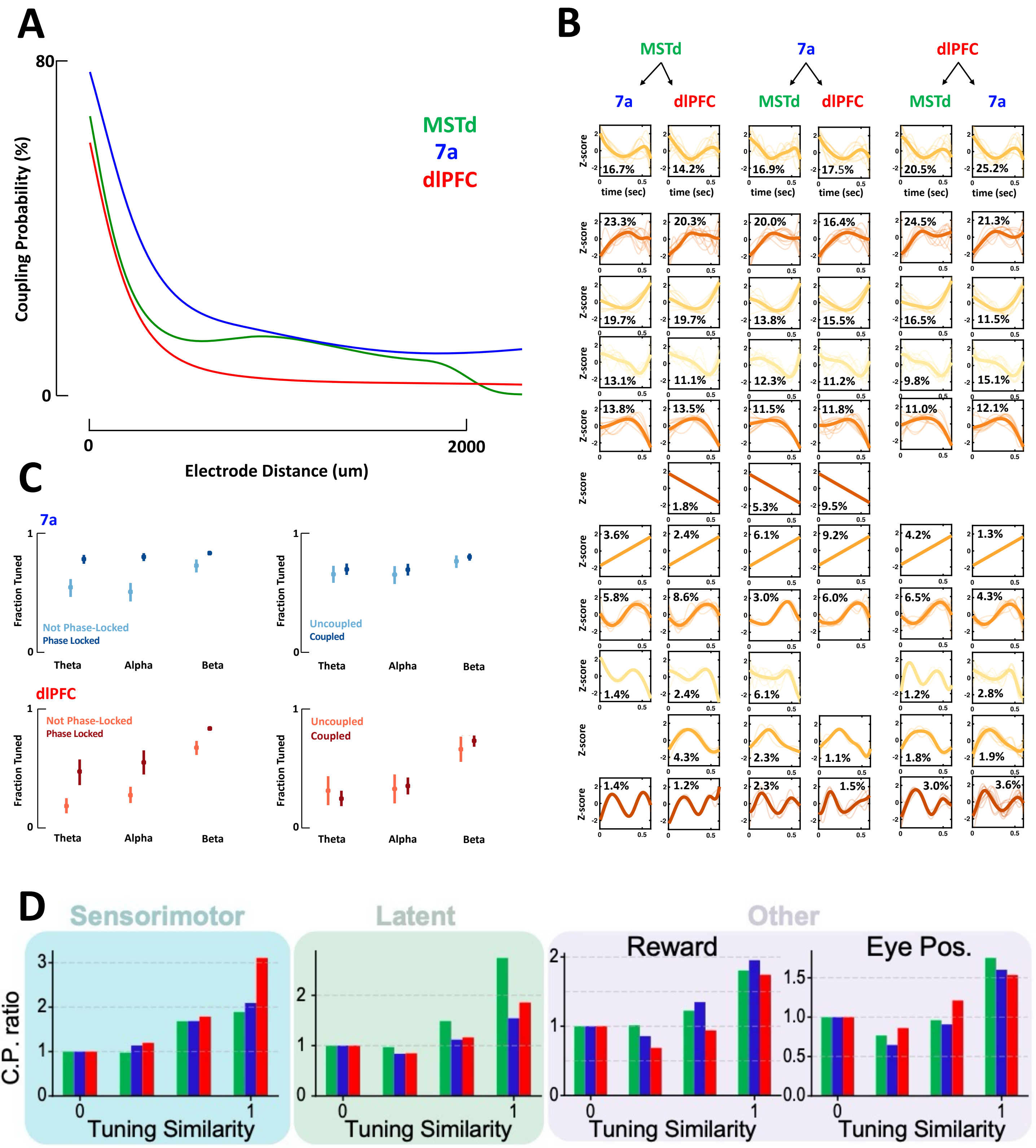

### Figure 5 Supplement 1

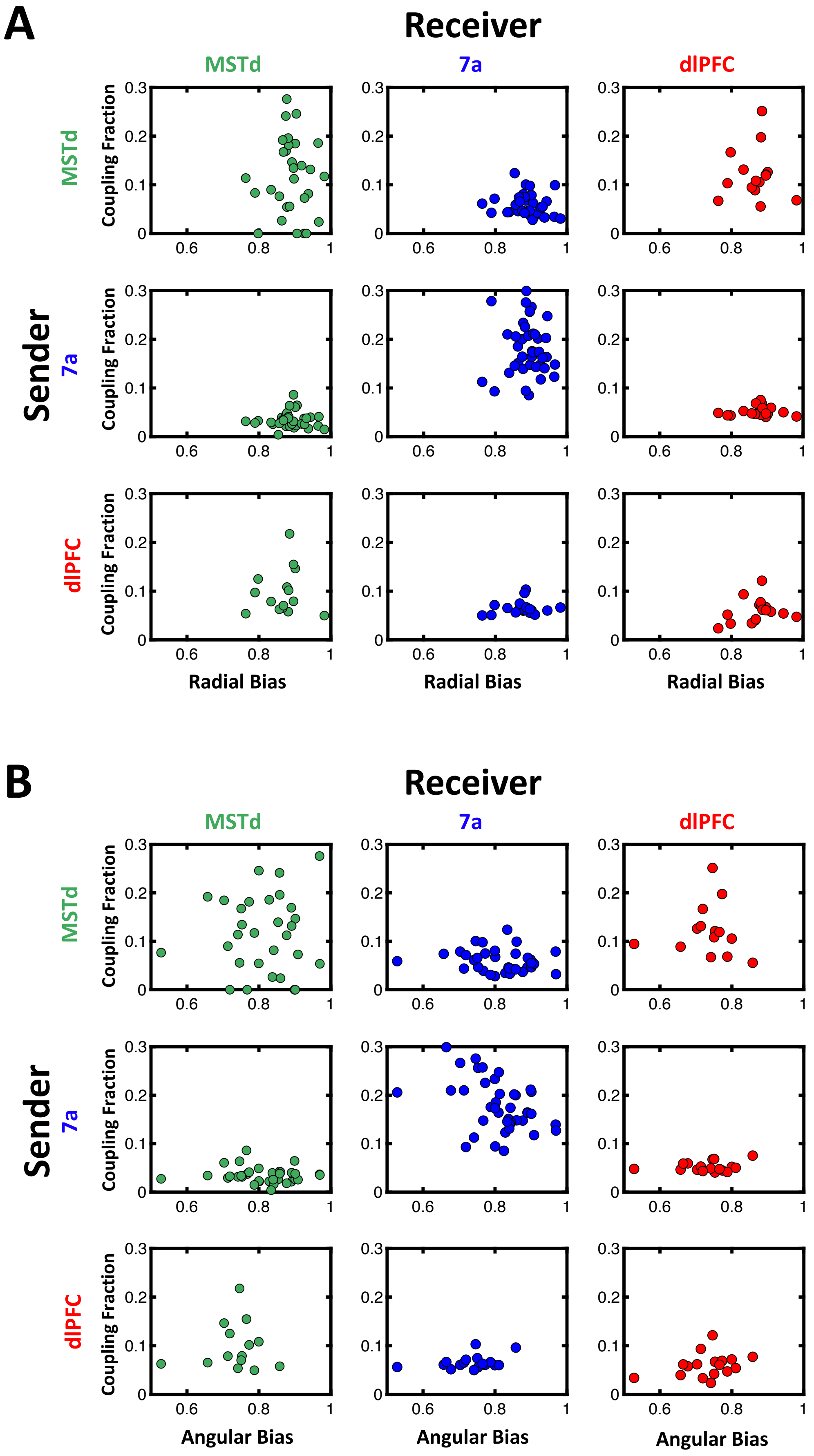
